# Multimodal molecular profiling of the metabolic penumbra in hyperacute stroke

**DOI:** 10.64898/2026.06.30.733797

**Authors:** Amin Mottahedin, Yvonne Couch, Paul Holloway, Philipp Mergenthaler, Philipp Boehm-Sturm, Moustafa Attar, Russell Foster, Andreas Dannhorn, Alastair M. Buchan

## Abstract

**Background:** The ischemic penumbra, a metabolically compromised yet potentially salvageable region surrounding the ischemic core, is a prime target for acute stroke intervention. Yet an objective molecular definition of the penumbra, particularly during the earliest stages of ischemia, remains lacking.

**Methods and Results:** We applied principal component analysis (PCA) followed by k-means clustering to high-resolution mass spectrometry imaging data covering multiple metabolic pathways to identify a metabolically defined penumbra in a mouse model of hyperacute stroke (45 min middle cerebral artery occlusion, MCAO). Targeted spatial metabolomic profiling by matrix-assisted laser desorption/ionization (MALDI) and desorption electrospray ionization (DESI) reveals a distinct penumbral metabolic profile, marked by relative preservation of high-energy phosphates, comparable lactate accumulation, and reduced succinate accumulation relative to the core. Spatial transcriptomics revealed selective induction of immediate-early genes, including *Npas4*, *Fos* and *Junb*, within the penumbra. Consistently, imaging mass cytometry shows enrichment of phospho-histone H3 (pHH3) within the penumbra, suggesting a chromatin-associated response potentially linked to immediate-early gene activation.

**Conclusion:** Together, these findings provide a multimodal molecular atlas of the hyperacute metabolically defined penumbra and reveal molecular features that facilitates its identification and inform future therapeutic strategies.

## Introduction

The aim of ischemic stroke treatment is to preserve viable brain tissue, primarily through the rapid restoration of cerebral blood flow. This salvageable tissue, known as the ischemic penumbra, has been characterised by severe hypoperfusion and electrical silence. Yet, unlike the irreversibly damaged infarct core, the reversible penumbra retains partial ionic homeostasis and metabolic viability^1–3^. Since its conceptualization, the penumbra has been central to the rationale behind reperfusion therapies, moving from a time-window to a tissue window^4^; however, its precise molecular definition remains incompletely resolved.

In both clinical and preclinical settings, advanced neuroimaging techniques have been instrumental in differentiating the dichotomous ischemic core from potentially salvageable tissue^5,6^. While these approaches have transformed the diagnosis of stroke and selection of patients for reperfusion therapies, they define the penumbra primarily through physiological and radiological characteristics rather than direct molecular measurements. Consequently, no objective molecular method currently exists to delineate the penumbra at cellular or subregional resolution, particularly during the hyperacute phase of ischemia when tissue fate remains unresolved and infarct boundaries are not yet clearly established. In experimental stroke studies, this often leads to penumbral tissue being operationally defined as peri-infarct regions surrounding the lesion rather than by objective molecular criteria. This lack of molecular definition has hampered mechanistic investigation of penumbral biology and limited the development of therapies specifically targeting tissue at risk during the narrow therapeutic window following stroke onset^7^.

Ischemic stroke triggers an immediate and profound metabolic crisis. Mass spectrometry-based metabolomic imaging has therefore emerged as a powerful tool to visualize and quantify spatially resolved metabolic alterations. Using DESI metabolic imaging in a mouse model of stroke, we recently demonstrated that mitochondrial succinate accumulates rapidly in ischemic tissue and is subsequently oxidized upon reperfusion, generating bursts of reactive oxygen species that contribute to tissue injury^8^. Pharmacological inhibition of succinate oxidation with malonate reduces infarct size and improves neurological outcome, underscoring the therapeutic value of metabolic interventions^8,9^. Yet, a major unresolved challenge is the lack of a robust method to discriminate metabolic responses in the penumbra from those in the core.

Here, we address this challenge by combining high-resolution mass spectrometry imaging with spatial transcriptomics and imaging mass cytometry. Rather than defining the penumbra according to anatomical location or a single molecular marker^10^, we used multidimensional metabolic signatures derived from a targeted panel of metabolites to delineate healthy tissue, penumbra and ischemic core in an unbiased manner. This approach enabled us to determine metabolic, transcriptional and protein-signalling changes in a spatially resolved way. Together, these data provide a unique molecular atlas of the hyperacute, metabolically-defined penumbra, and establish a framework for identifying molecular features associated with tissue salvageability.

## Material and Methods

### Animals

Adult male C57Bl6J mice (Charles River, UK) at 12-14 weeks of age were housed under 12:12 hour light-dark cycle (lights on at 7 AM and lights off at 7 PM) with *ad libitum* access to food and water. All procedures were carried out in accord with the UK Animals (Scientific Procedures) Act (1986) and licensed protocols were approved by local committees (LERP and ACER, University of Oxford) and carried out under license number PP7444704.

### Middle Cerebral Artery Occlusion (MCAO)

The MCAO surgeries were performed at zeitgeber time 3 (ZT3, 10 AM). Mice were anesthetised by isoflurane (3% for induction, 1.5-2% maintenance), common carotid artery and external carotid artery were exposed and ligated, a silicone-tipped filament (0.22mm diameter, 2-3 mm coating length. Doccol, USA) was inserted into internal carotid artery and advanced to occlude MCA. The occlusion was confirmed when at least 70% drop observed in cerebral flood flow as measured by doppler flowmetry (Oxford Optronix, UK). All three animals achieved >70% reduction in cerebral blood flow, and no animals were excluded. After 45 min of MCAO under anaesthesia, mice were sacrificed by cervical dislocation, and brain was rapidly dissected and snap-frozen in liquid nitrogen-cold isopentane.

### Mass spectrometry imaging (MSI)

Brain tissues were embedded in a HPMC/PVP hydrogel according to a published protocol to generate multi-tissue blocks^11^. Blocks were cryo-sectioned at 10 μm thickness on a cryostat (Leica Biosystems, Germany). Sections were thaw-mounted onto SuperFrost slides (Thermo Scientific, Germany) for DESI-MSI and ITO-coated slides for MALDI-MSI. Serial sections were collected across eight rostrocaudal tiers spanning approximately Bregma −3 mm to +5 mm at 1 mm intervals. At each interval 25 sections were cut and collected. Six tiers were selected for MSI analysis to ensure representation of metabolic heterogeneity across the lesion. Sections were air-dried, vacuum-sealed, and stored at −80 °C until analysis.

DESI-MSI was performed on a Q Exactive Plus mass spectrometer (Thermo Scientific, UK) equipped with an automated DESI ion source (Prosolia Inc., USA). Data were acquired in negative ion mode over an m/z range of 80–1000 at a nominal mass resolution of 70,000. The injection time was 150 ms, yielding a scan rate of 3.8 pixels s⁻¹. A custom-built DESI sprayer delivered 95% methanol/ 5% water at 1.5 μl min⁻¹ and was nebulized with nitrogen at 6 bar back pressure. The spatial resolution was 100 μm.

MALDI-MSI was performed on a RapifleX instrument (Bruker Daltonics, Bremen, Germany) using 9-aminoacridine (9-AA) as the matrix. Spectra were acquired in negative ion mode across m/z 100–900 with a spatial resolution of 65 μm. A total of 500 laser shots per pixel were accumulated to generate the final spectra.

Data processing and analysis were conducted in SCiLS Lab (v2024, Bruker Daltonics, Germany). Brain tissue was delineated using a partial least squares (PLS) classifier trained on manual annotations. Ischemic regions were identified through principal component analysis (PCA), followed by bisecting k-means clustering based on the most discriminative PCA loadings. Adjacent sections collected for histology were stained with hematoxylin and eosin (H&E) and imaged at 20× magnification using an Aperio scanner (Leica Biosystems, Germany). Other adjacent sections were used for spatial transcriptomics and imaging mass cytometry.

### GeoMx spatial transcriptomics

Spatial transcriptomic profiling was performed using the GeoMx Digital Spatial Profiler (NanoString Technologies) and the Mouse Whole Transcriptome Atlas (WTA). Fresh-frozen brain sections adjacent to those used for mass spectrometry imaging were processed according to the manufacturer’s protocol. Regions of interest (ROIs) corresponding to healthy tissue, metabolically defined penumbra and core were selected on the basis of adjacent MSI-derived segmentation maps. Libraries were generated using the GeoMx WTA workflow and sequenced by Azenta Life Sciences (GENEWIZ) on an Illumina NovaSeq platform using 2 × 150 bp paired-end sequencing.

Raw transcript counts were processed using the GeoMx analysis suite. Gene expression values were normalized using the geometric mean of housekeeping genes (*Gapdh, Actb, Hprt, Eef2, Ywhaz* and *Sdha*). Differential expression analysis was performed in R (v4.5.1) using the limma package. Normalized counts were transformed as log2(count + 1), linear models were fitted for each comparison, and statistical significance was assessed using empirical Bayes moderation. P values were adjusted for multiple testing using the Benjamini–Hochberg procedure. Genes with an adjusted *P* value < 0.05 and an absolute log2 fold change > 1 were considered differentially expressed.

### Imaging Mass Cytometry

Imaging mass cytometry (IMC) was performed on brain sections adjacent to those used for mass spectrometry imaging. Antibodies used in the IMC panel are listed in Supplementary Table 1. Antibodies were conjugated to metal isotopes using the Maxpar Antibody Labeling Kit (Fluidigm) according to the manufacturer’s instructions. Tissue sections were fixed in 4% paraformaldehyde in PBS for 10 min, washed three times in PBS and permeabilized using Triton X-100 (1:1,000 dilution in casein solution). Following three additional PBS washes, sections were blocked in casein solution (Thermo Fisher Scientific) for 30 min at room temperature. Metal-conjugated antibodies were diluted in casein solution and incubated overnight at 4 °C.

Following primary antibody incubation, sections were washed three times in PBS and nuclei were stained with DNA Intercalator-Iridium (Fluidigm; 1:400 dilution in PBS) for 30 min. Slides were subsequently washed three times in PBS, briefly rinsed in deionized water and air-dried at room temperature.

Regions of interest were selected using corresponding MSI-derived segmentation maps to identify healthy tissue, metabolically defined penumbra and core. IMC data were acquired using a Hyperion Imaging System (Fluidigm) with an ablation energy of 4 dB and an ablation frequency of 200 Hz. Images were visualized using MCD Viewer (v1.0, Fluidigm) applying same thresholds for all ROIs.

### Statistical analysis

Statistical analyses of metabolomic imaging data were performed using GraphPad Prism (v11.0.0; GraphPad Software). Relative metabolite abundances were compared across healthy tissue, metabolically defined penumbra and core using one-way analysis of variance (ANOVA) followed by Tukey’s multiple-comparisons test. Data are presented as mean ± SD. Exact *P* values are reported in the figures, and statistical significance was defined as *P* < 0.05.

## Results

### Mass spectrometry imaging identifies a metabolically distinct ischemic penumbra

To define the molecular landscape of hyperacute ischemic tissue and identify the metabolic penumbra in an unbiased manner, we performed high-resolution mass spectrometry imaging (MSI) on serial brain cryosections from mice subjected to 45 min middle cerebral artery occlusion (MCAO) (Fig. 1A). Principal component analysis (PCA) of MSI datasets revealed a continuous metabolic trajectory spanning healthy and ischemic tissue, with the second principal component capturing regional heterogeneity within the ischemic hemisphere (Fig. 1B). Projection of PCA loadings onto tissue sections identified three spatially distinct metabolic compartments corresponding to healthy tissue, a peripheral penumbra, and a central ischemic core, while no clearly demarcated lesion was evident on H&E staining at this hyperacute stage (Fig. 1C). Data-driven segmentation based on PCA-derived features consistently delineated these regions across serial sections spanning the rostrocaudal extent of the lesion (Fig. 1D). Metabolic classification was supported by characteristic distributions of lactate and succinate. Lactate accumulated throughout the ischemic region, whereas succinate enrichment was concentrated predominantly within the core region (Fig. 1C). Quantification of segmented volumes demonstrated that healthy tissue represented 64.1±2.4% (mean±SD) of total brain volume, while metabolically defined penumbra and core occupied 15.9±2% and 20±3.7%, respectively (Fig. 1E). These findings establish MSI-based metabolic segmentation as a robust approach for defining the metabolic penumbra during the hyperacute phase of stroke.

**Figure 1.**
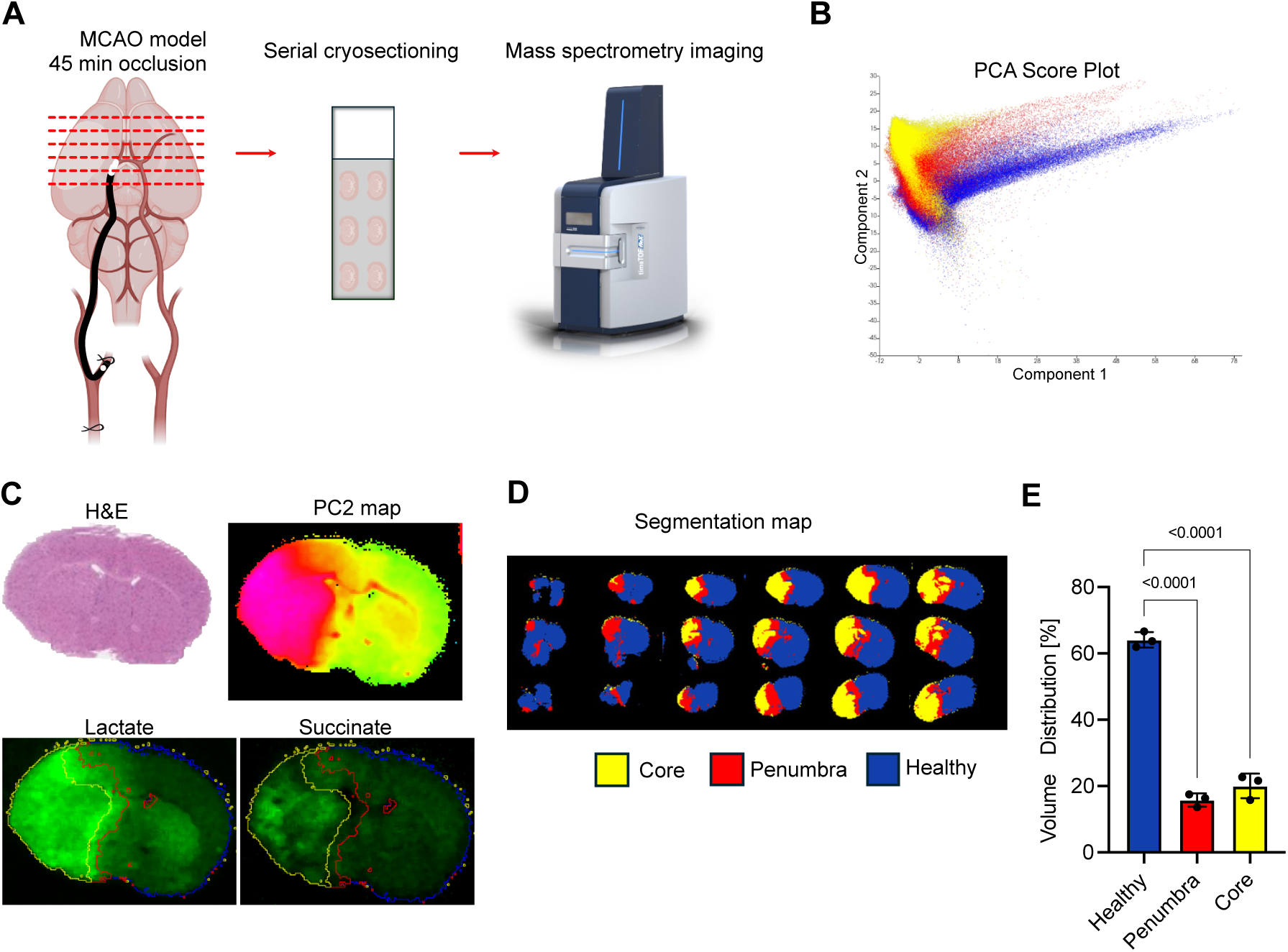
| Mass spectrometry imaging identifies a metabolically defined ischemic penumbra. **A**, Experimental workflow. Mice were subjected to 45 min middle cerebral artery occlusion (MCAO; modified from BioRender.com), followed by brain collection, serial cryosectioning and mass spectrometry imaging (MSI). Adjacent sections were reserved for complementary spatial analyses. **B,** Principal component analysis (PCA) score plot of MSI data demonstrating separation of metabolic states within the ischemic hemisphere. **C,** Representative PCA component 2 (PC2) map identifying distinct metabolic regions and corresponding H&E staining on adjacent section. Spatial distributions of lactate and succinate are shown with yellow, red, and blue lines demarcating core, penumbra and healthy regions, respectively. **D,** Data-driven segmentation of serial brain sections based on PCA-derived metabolic features. **E,** Quantification of tissue volume distribution assigned to healthy tissue, penumbra and core following segmentation. Data are presented as mean ± SD. Statistical significance was determined by one-way ANOVA with multiple-comparison correction. Exact *P* values are indicated in the figure. N=3.

### The metabolically defined penumbra exhibits a distinct profile of glycolytic and tricarboxylic acid cycle metabolism

We next examined spatial changes in central carbon metabolism across healthy tissue, metabolically defined penumbra and core (Fig. 2A). Glycolytic intermediates demonstrated a progressive shift consistent with impaired oxidative metabolism. Glucose abundance was reduced in the core compared with both healthy tissue and penumbra, whereas penumbral glucose levels remained comparable to healthy tissue (Fig. 2B). Pyruvate was significantly decreased in both ischemic compartments relative to healthy tissue, with the greatest reduction observed in the core (Fig. 2C). In contrast, lactate accumulated markedly in both penumbra and core, reaching similar levels in the two ischemic regions and indicating extensive engagement of glycolysis (Fig. 2D).

**Figure 2.**
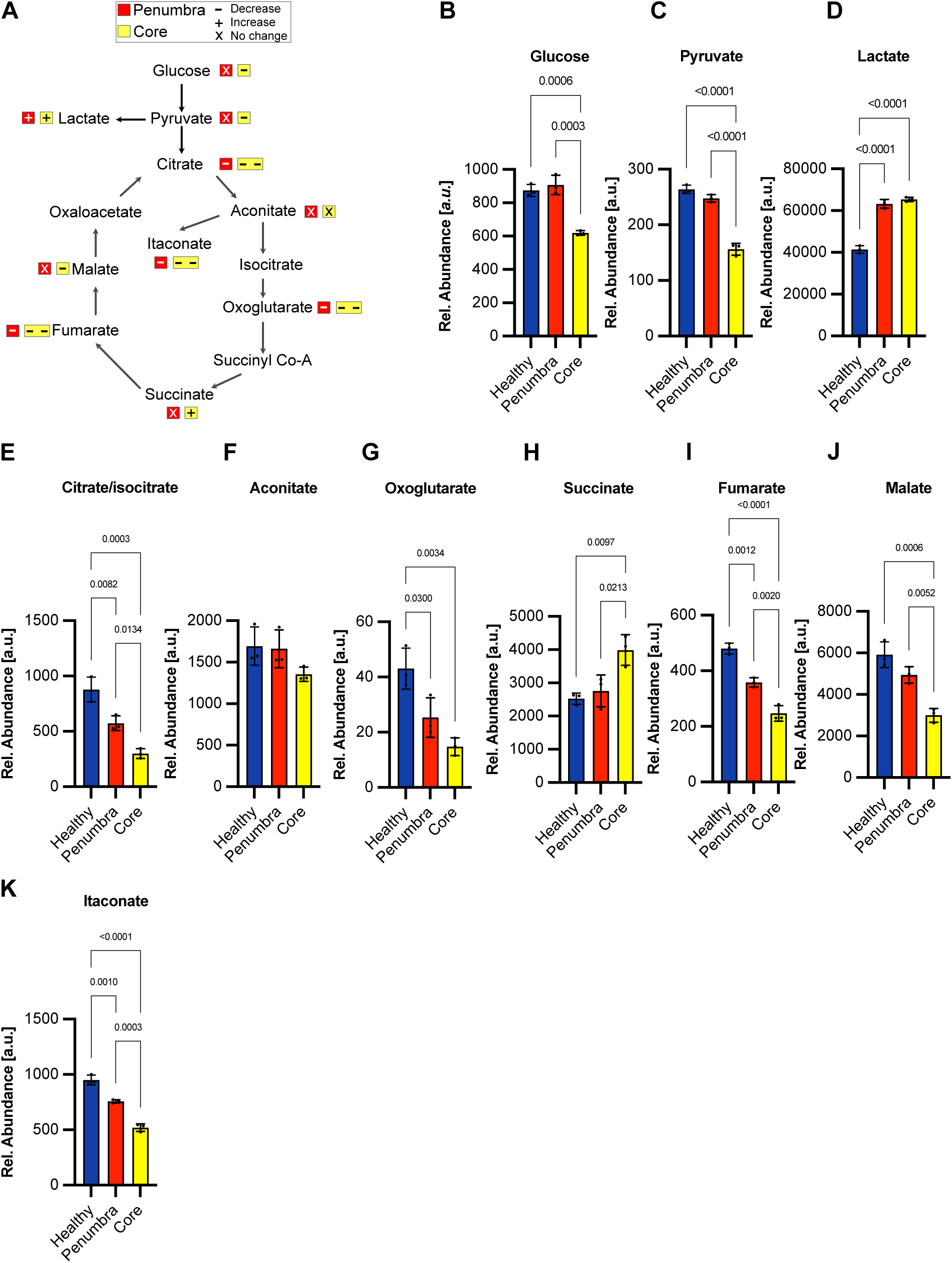
| Spatial remodelling of glycolytic and tricarboxylic acid cycle metabolism in the ischemic region. **A**, Schematic summary of metabolic changes in glycolysis and the tricarboxylic acid (TCA) cycle within penumbra and core relative to healthy tissue. **B-K,** Relative abundance of glycolytic and TCA-cycle metabolites measured by MSI in healthy tissue, penumbra and ischemic core. **B,** Glucose. **C,** Pyruvate. **D,** Lactate. **E,** Citrate/isocitrate. **F,** Aconitate. **G,** Oxoglutarate. **H,** Succinate. **I,** Fumarate. **J,** Malate. **K,** Itaconate. Data are presented as mean ± SD. Statistical significance was determined by one-way ANOVA with multiple-comparison correction. Exact *P* values are shown in the figure. N=3.

Analysis of tricarboxylic acid (TCA) cycle intermediates revealed pronounced metabolic divergence between penumbra and core. Citrate/isocitrate abundance decreased progressively from healthy tissue to penumbra and further to the core (Fig. 2E), whereas aconitate remained largely unchanged across regions (Fig. 2F). Oxoglutarate was reduced in both ischemic compartments, with the lowest levels observed in the core (Fig. 2G). In contrast, succinate accumulated selectively within ischemic tissue and reached its highest abundance in the core (Fig. 2H). Downstream metabolites fumarate and malate showed the opposite pattern, decreasing in penumbra and exhibiting further reductions in the core (Fig. 2I,J). The immunometabolite itaconate was also significantly reduced in ischemic tissue, with the greatest depletion observed in the core (Fig. 2K). Collectively, these findings indicate that although both penumbra and core experience substantial metabolic stress, the penumbra retains a partially preserved TCA cycle state characterized by lower succinate accumulation and greater maintenance of downstream intermediates compared with the metabolically collapsed core.

### Relative preservation of adenine nucleotide metabolism distinguishes the metabolically defined penumbra from the core

Because failure of cellular energy metabolism is a defining feature of ischemic injury, we next assessed the spatial distribution of adenine nucleotides and their degradation products (Fig. 3A). ATP levels were significantly reduced in both ischemic compartments compared with healthy tissue, with the most profound depletion occurring in the core (Fig. 3B). However, ADP and AMP level were significantly reduced only in the ischemic core (Fig. 3C-D). Consistently, adenosine, a key product of ATP catabolism, increased significantly in the core only (Fig. 3E). Further downstream metabolites exhibited marked regional differences. Adenine (Fig. 3F), inosine (Fig. 3G) and xanthine (Fig. 3H) accumulated progressively across the ischemic gradient and reached their highest levels in the core.

**Figure 3.**
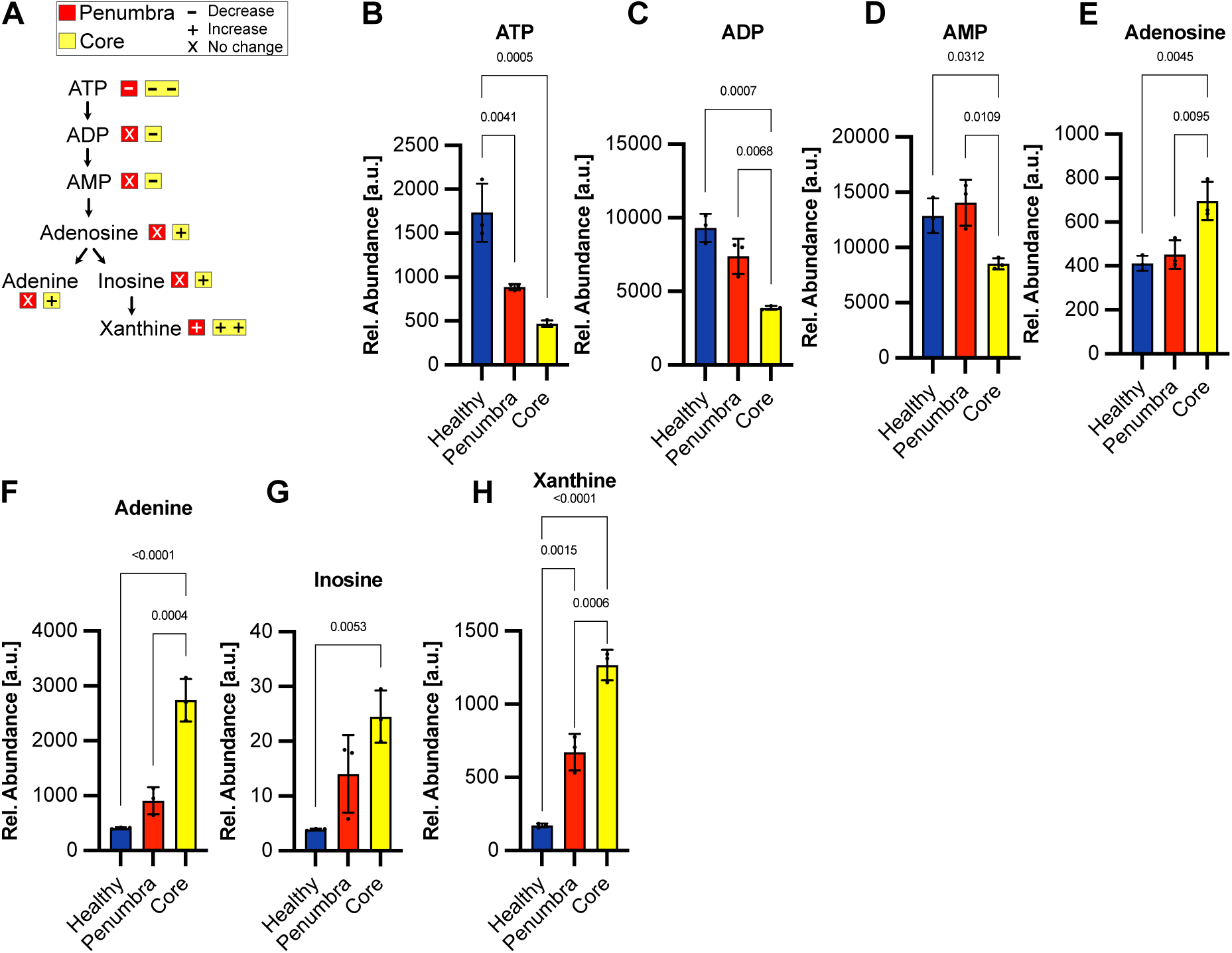
| Differential regulation of adenine nucleotide metabolism across healthy tissue, penumbra and ischemic core. **A**, Schematic overview of adenine nucleotide metabolism highlighting metabolites altered within penumbra and core relative to healthy tissue. **B–H,** Relative abundance of adenine nucleotides and degradation products measured by MSI. **B,** ATP. **C,** ADP. **D,** AMP. **E,** Adenosine. **F,** Adenine. **G,** Inosine. **H,** Xanthine. Data are presented as mean ± SD. Statistical significance was determined by one-way ANOVA with multiple-comparison correction. Exact *P* values are indicated in the figure. N=3.

Together, these data demonstrate progressive degradation of the adenine nucleotide pool across the ischemic lesion. Compared with the core, the penumbra retains higher ATP and ADP levels and exhibits less accumulation of purine catabolites, consistent with partial preservation of energy metabolism within metabolically salvageable tissue.

### Fatty acid metabolism is differentially altered across the ischemic region

To investigate whether fatty acid metabolism differs between the metabolically defined penumbra and core, we quantified a panel of fatty acid and oxidized fatty acid species by MSI (Fig. 4). Oxidized FA(6:0)+O was selectively reduced in the core relative to both healthy tissue and penumbra, whereas penumbral levels remained comparable to healthy tissue (Fig. 4A). FA(8:1)+2O was decreased in both penumbra and core compared with healthy tissue, indicating an ischemia-associated reduction that was not specific to either compartment (Fig. 4B). Similarly, the unsaturated long-chain fatty acids FA(16:1) and FA(18:2) were significantly reduced in the core but remained preserved within the penumbra (Fig. 4C,D). In contrast, arachidonic acid (FA(20:4)) exhibited a progressive increase across the ischemic gradient, with levels rising from healthy tissue to penumbra and reaching their highest abundance within the core (Fig. 4E). The oxidized derivative FA(20:4)+O displayed a distinct pattern, showing a trend towards reduced abundance in the penumbra while accumulating significantly within the core (Fig. 4F). The remaining fatty acid species analysed did not exhibit significant differences between the ischemic compartments (Supplementary Fig. S1). Together, these findings reveal compartment-specific remodelling of fatty acid metabolism during hyperacute ischemia, with relative preservation of several fatty acid species within the metabolically defined penumbra and more profound alterations within the ischemic core.

**Figure 4.**
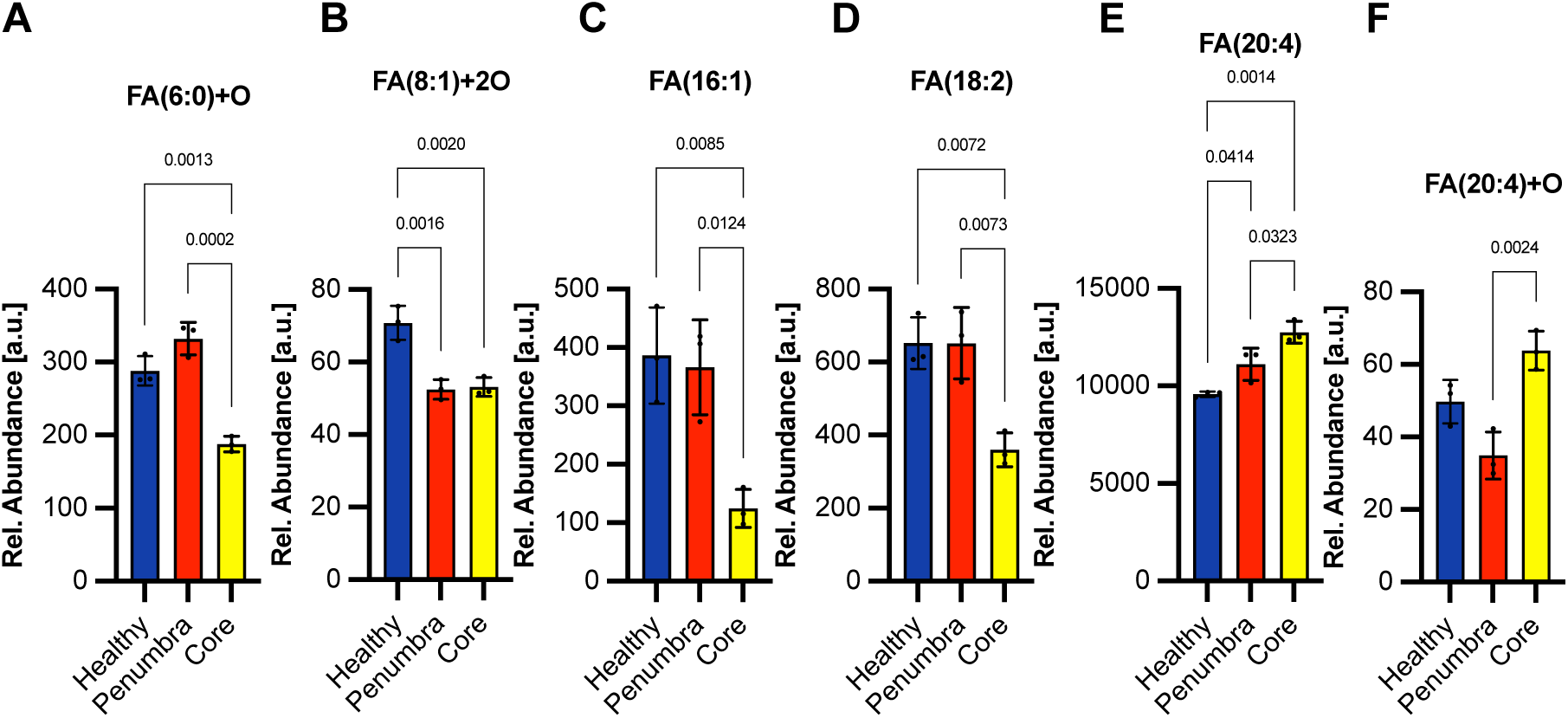
| Spatial alterations in fatty acid metabolism during hyperacute cerebral ischemia. **A–F**, Relative abundance of fatty acid and oxidized fatty acid species measured by MSI in healthy tissue, penumbra and ischemic core. **A,** FA(6:0)+O. **B,** FA(8:1)+2O. **C,** FA(16:1). **D,** FA(18:2). **E,** FA(20:4) (arachidonic acid). **F,** FA(20:4)+O. Data are presented as mean ± SD. Statistical significance was determined by one-way ANOVA with multiple-comparison correction. Exact *P* values are shown in the figure. N=3.

### The metabolically defined penumbra exhibits a distinct immediate-early gene transcriptional program

To determine whether the metabolically defined penumbra represents a transcriptionally distinct tissue compartment, we performed GeoMx spatial transcriptomics on regions of interest (ROIs) selected from healthy tissue, metabolically defined penumbra and core identified by MSI-based segmentation (Fig. 5A). Comparison of penumbral ROIs with contralateral healthy tissue revealed a discrete but robust transcriptional response characterized by selective upregulation of immediate-early genes (IEGs) (Fig. 5B, Supplementary Table 2). Among the most strongly induced transcripts were *Npas4*, *Fosb*, *Fos*, *Nr4a1*, *Junb*, *Dusp1*, *Btg2*, *Arl4d* and *Arc*, all of which are established immediate-early genes (IEGs) involved in rapid cellular adaptation to stress and neuronal stimulation^12^.

**Figure 5.**
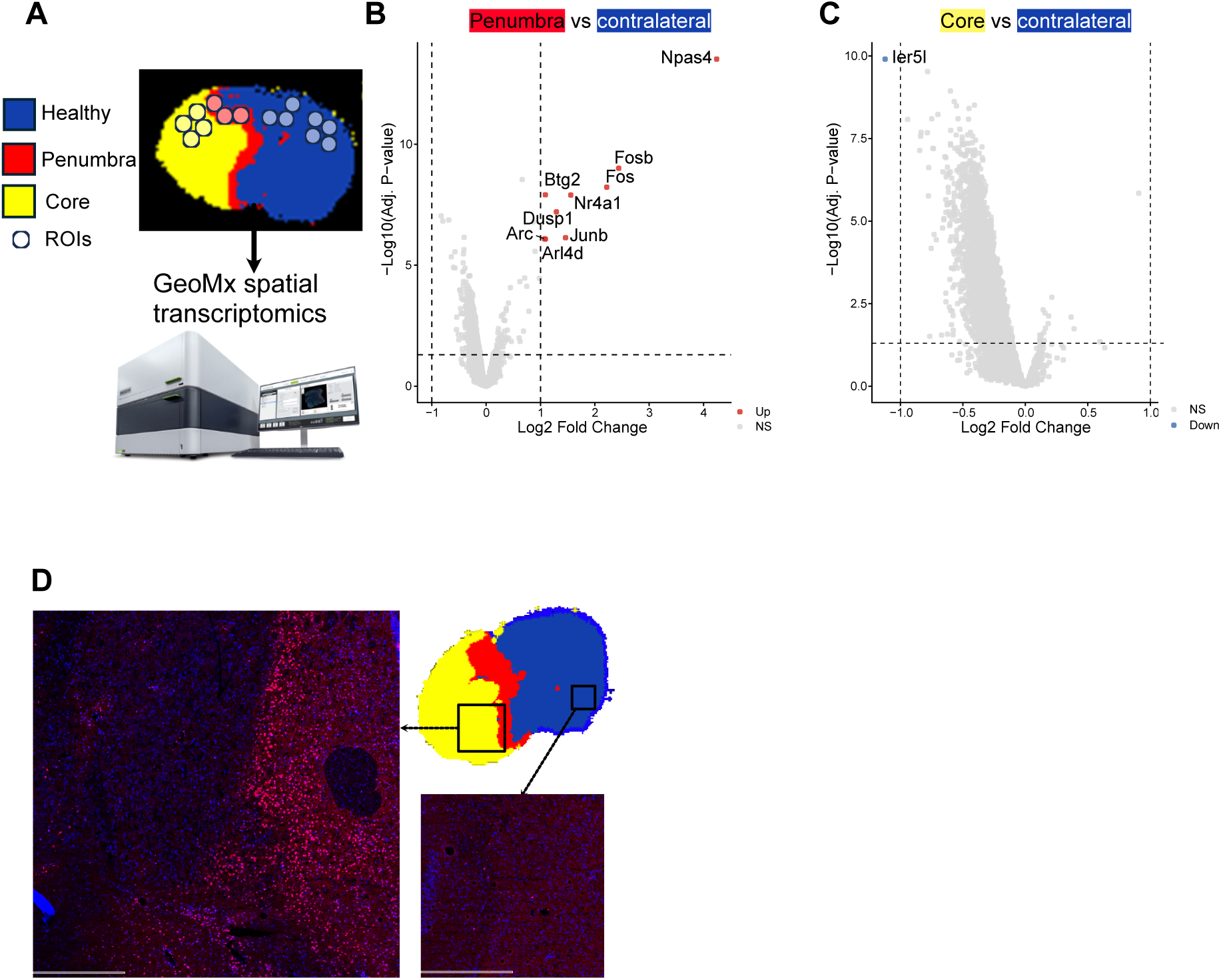
| Spatial transcriptomics identifies a penumbra-specific immediate-early gene response associated with phospho-histone H3 enrichment. **A**, Experimental workflow showing selection of regions of interest (ROIs) from MSI-defined healthy tissue (blue), penumbra (red) and ischemic core (yellow) for GeoMx spatial transcriptomic analysis. **B**, Volcano plot showing differential gene expression between penumbra and contralateral healthy tissue. Red circles represent differentially upregulated genes with adjusted p<0.05 and log2 fold change>1. **C**, Volcano plot showing differential gene expression between ischemic core and contralateral healthy tissue. Blue circles represent differentially downregulated genes with adjusted p<0.05 and log2 fold change>1.**D**, Representative phospho-histone H3 (pHH3, red) and nuclear (blue, 193Ir) staining demonstrating enrichment at the metabolically defined penumbra. ROI in ipsilateral hemisphere is 2X2mm and in contralateral hemisphere is 1X1mm. Scale bar = 500µM.

In contrast, comparison of the ischemic core with contralateral tissue identified only a single differentially downregulated gene, *Ier5l*, with a log2 fold change <-1 (Fig. 5C). Rather than exhibiting induction of distinct transcriptional programs, the core was characterized by a slight reduction in the abundance of hundreds of genes (Supplementary Table 2), consistent with severe metabolic failure and suggesting global suppression of transcriptional activity and/or potentially extensive loss of RNA integrity.

To determine whether the metabolically defined penumbra also exhibits a distinct protein-signalling state, we performed imaging mass cytometry (IMC) on adjacent tissue sections using a multiplexed antibody panel encompassing neuronal, vascular, immune, metabolic and phospho-signalling phenotypic and architectural markers (Supplementary Table 1). Regions analysed by IMC were aligned with MSI-derived maps to enable direct comparison between healthy tissue, penumbra and ischemic core. Across the panel, some markers displayed attenuated signal within the core (e.g. b-catenin), consistent with severe tissue injury (Supplementary Fig. S2). In contrast, chromatin-associated signal, phospho-histone H3 (pHH3) exhibited a striking spatial pattern. pHH3 signal was enriched within the metabolically defined penumbra and was largely absent from both healthy tissue and the ischemic core. Its spatial distribution closely mirrored that of the penumbra-specific immediate-early gene response. Together, these findings indicate that the metabolically defined penumbra is associated with a distinct transcriptional program as well as a phospho-signalling state, indicative of chromatin remodelling, that is not observed in either healthy tissue or the ischemic core.

## Discussion

We used data-driven segmentation based on multiple metabolic pathways, including glycolysis, TCA-cycle metabolism, adenine nucleotides and lipid species, allowing identification of a metabolically defined penumbra independent of anatomical location or any single biomarker. The ischemic penumbra is the principal target of acute stroke therapies, yet its molecular identity remains incompletely understood^13,14^. In experimental mouse MCAO models, the ischemic penumbra is a transient phenomenon that evolves rapidly over the first several hours after stroke onset. Without reperfusion, the penumbra progressively converts into irreversibly damaged tissue, with substantial loss of salvageable tissue generally occurring within 3–6 hours. As such, understanding the molecular landscape early in this process will be key to defining potential therapeutic targets in future studies.

Mass spectrometry imaging has been used previously to investigate metabolic heterogeneity in ischemic stroke, but mainly in later phases of injury, at a time when the infarct is largely established^15^. One notable exception is the work of Hattori and colleagues, who examined adenine nucleotide metabolism during MCA occlusion and reported relative preservation of ATP within peri-core tissue^16^. However, tissue classification was based primarily on adenylate distributions, with peri-core regions assumed to represent penumbra.

Carbon metabolism is among the earliest cellular processes disrupted following cerebral ischaemia, reflecting the transition from aerobic respiration to alternative metabolic pathways in response to energy failure. Indeed, here we found that one of the features of the metabolically defined penumbra was changes in central carbon metabolism. Lactate accumulated to similar levels in both penumbra and core, indicating high glycolytic activity throughout the ischemic tissue. In contrast, TCA-cycle metabolites revealed substantial metabolic divergence between the two compartments.

We previously demonstrated that succinate accumulates rapidly during cerebral ischemia and its oxidation at reperfusion is a main driver of ischemia-reperfusion injury^8,9^. Gruszczyk and colleagues showed that both hypoxia and anoxia induce succinate accumulation in cardiomyocytes, but that the magnitude of accumulation is several-fold greater under anoxic conditions, suggesting that the accumulation of succinate is highly sensitive to oxygen availability.^17^. Here, succinate accumulated predominantly within the core, whereas downstream metabolites such as fumarate and malate were relatively preserved within the penumbra. This lower succinate abundance in the presence of preserved downstream TCA-cycle intermediates, as well as maintenance of ATP and ADP in the penumbra suggests a gradient of ischemic severity. These data demonstrate that where the core undergoes profound metabolic collapse, penumbral tissue likely retains sufficient metabolic capacity to sustain partial oxidative metabolism.

In addition to alterations in central carbon metabolism, we observed pronounced differences in lipid metabolism between penumbra and core. Disturbances in fatty-acid homeostasis occur rapidly following cerebral ischemia, with increases in free fatty acids and arachidonic acid have previously been reported within minutes of vessel occlusion^18–20^. However, these studies relied predominantly on bulk tissue extraction or without discriminating between metabolically distinct tissue states within the ischemic region^21^. By defining the penumbra through unbiased metabolic segmentation, our study reveals that lipid remodelling is highly compartmentalized during hyperacute ischemia, with preservation of several fatty-acid species within the penumbra and preferential accumulation of arachidonic acid within the core. Given the established roles of arachidonic acid metabolism in oxidative stress, inflammation and membrane degradation^22^, these findings suggest that lipid-mediated injury pathways are activated most strongly within irreversibly damaged tissue. Conversely, preservation of several lipid species within the penumbra may reflect maintenance of membrane integrity and lipid homeostasis despite ongoing ischemic stress.

Spatial transcriptomics revealed that the metabolically defined penumbra remained transcriptionally active despite severe metabolic stress. It identified robust induction of immediate-early genes including Npas4, Fos, Fosb, Junb, Arc, Nr4a1 and Dusp1, whereas the ischemic core exhibited broad but modest transcriptional suppression or degradation. Immediate-early gene (IEG) induction following cerebral ischemia has been recognized for more than three decades, with studies reporting activation of c-fos, c-jun, junB and NGFI family genes within hours of focal or global ischemia^12^. However, again, these studies generally defined penumbral tissue anatomically, based on proximity to the established lesion, rather than functionally. Consequently, the relationship between induction of IEGs and the molecular state of hyperacute salvageable tissue has remained unclear. Our findings demonstrate that IEG activation is a hallmark of the hyperacute metabolic penumbra. Notably, Npas4 was among the most strongly induced transcripts. Previous studies have identified Npas4 as a rapidly induced activity-dependent transcription factor with neuroprotective functions in experimental cerebral ischemia^23^. Its prominent induction within the metabolically defined penumbra suggests that endogenous adaptive stress-response pathways remain active within this compartment despite severe metabolic compromise.

The transcriptionally active nature of the penumbra was further supported by imaging mass cytometry, which identified phospho-histone H3 (pHH3) enrichment within this compartment. Histone H3 phosphorylation has been linked to both chromatin remodelling and cell-cycle regulation, depending on the biological context^24^. Previous studies reported pHH3-positive cells within anatomically defined penumbral regions 24-72 h after experimental stroke, where pHH3 was primarily interpreted as a marker of cell-cycle activation and neurogenesis^25^. In contrast, we observed pHH3 enrichment within a metabolically defined penumbra during the hyperacute phase of ischemia. Notably, the spatial distribution of pHH3 closely mirrored the penumbra-specific immediate-early gene programme identified by spatial transcriptomics. Although the functional relationship between these observations remains to be determined, the findings suggest that pHH3 marks an early molecular response associated with metabolically salvageable tissue.

Together, these findings provide a multimodal molecular atlas of the hyperacute metabolic penumbra and establish an objective molecular framework for identifying tissue at risk at the earliest stages of ischemic injury. More broadly, they highlight the molecular complexity of the penumbra, extending the concept beyond a singular anatomically defined region towards a coordinated metabolic and transcriptional tissue state. The molecular features identified here may also provide a foundation for developing simpler approaches to define and target the penumbra.

### Limitations

Several limitations should be acknowledged. First, the number of animals analysed was modest. However, this is consistent with our (and others’) previous mass spectrometry imaging studies in experimental stroke, where metabolic signatures showed low variability across biological replicates^8,15^. Furthermore, the key findings reported here were independently supported by three orthogonal modalities, mass spectrometry metabolic imaging, spatial transcriptomics and imaging mass cytometry, providing confidence in the robustness of the observed molecular patterns. Second, the study was performed exclusively in young male mice and therefore does not address the influence of sex, ageing or common comorbidities on penumbral biology.

Although our unbiased metabolic segmentation identified spatially distinct regions corresponding to healthy tissue, penumbra and ischemic core, these classifications should be interpreted as an analytical framework rather than discrete biological entities. Ischemic injury evolves along continuous gradients of perfusion, oxygen availability, metabolic dysfunction and cellular viability, and recent work has highlighted that the traditional core-penumbra model may not fully capture this heterogeneity^25,26^. Consistent with this concept, our data revealed gradual spatial transitions in metabolites including succinate, adenine nucleotides and fatty acids, suggesting that metabolic states change progressively across the lesion rather than at fixed boundaries. Rather than diminishing the utility of our approach, these findings demonstrate that spatial metabolomics can resolve the molecular landscape of ischemic injury, whilst also providing an objective framework for interrogating tissue with salvage potential. Future studies combining higher spatial resolution with additional timepoints may further delineate intermediate metabolic states and refine our definition of the ischemic penumbra.

## Author contributions

AM conceived and designed the study. AM, AD, PM, PBS, and AMB designed the experiments. AM, AD, YC, and MA performed the experiments. AM and AD analysed the data. AM, YC, PH, PM, and AMB interpreted the data. AMB and RF acquired funding, provided infrastructure and resources, and supervised the project. AM drafted the manuscript. All authors contributed to manuscript revision, editing and approved the final version.

## Statements and Declarations

AMB is a co-founder of Brainomix, the other authors declare no competing interests.

---

**Supplementary Figure S1.**
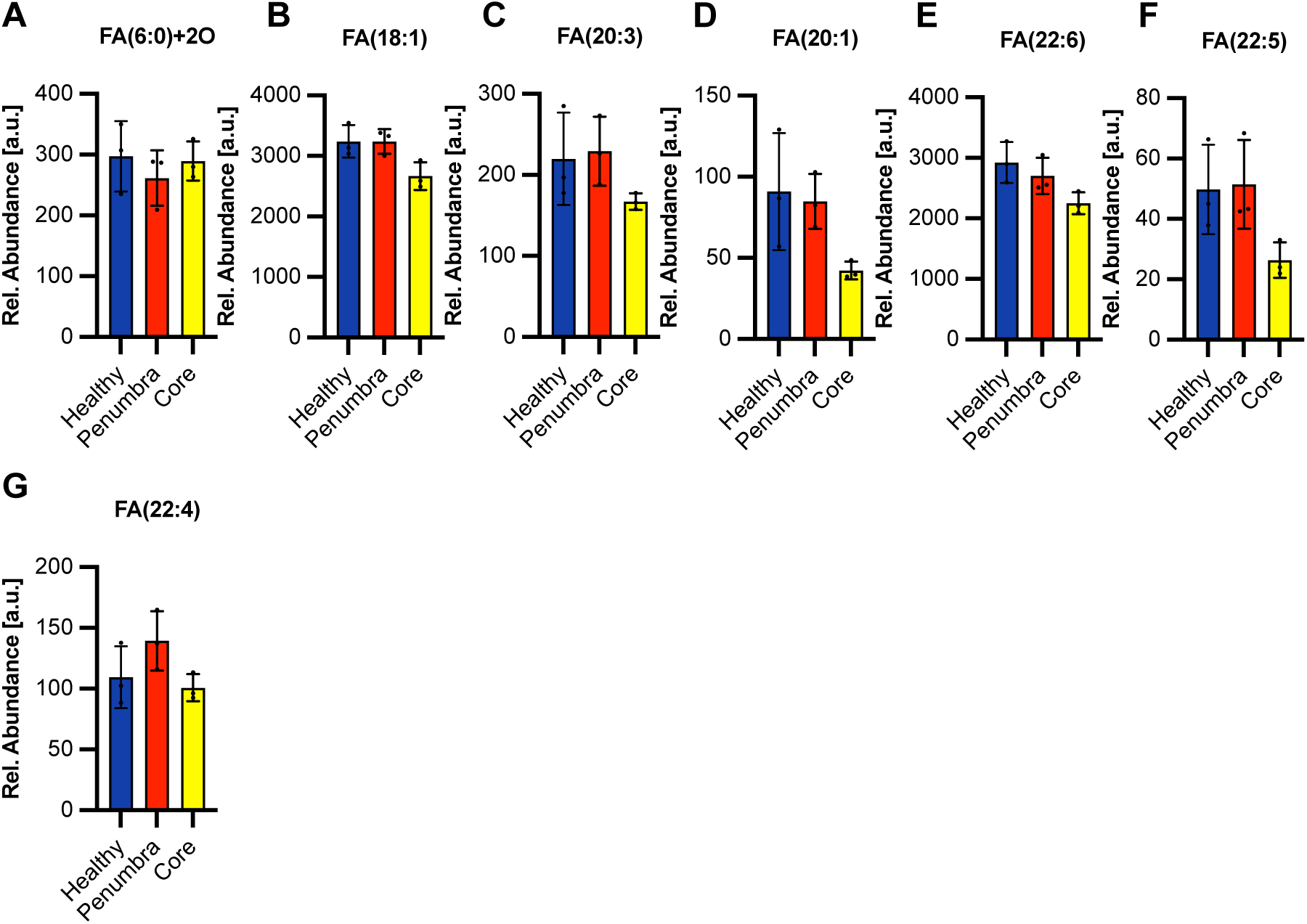
| Additional fatty acid species quantified by mass spectrometry imaging. Relative abundance of additional fatty acid and oxidized fatty acid species measured by MSI in healthy tissue, penumbra and ischemic core. (A) FA(6:0)+2O, (B) FA(18:1), (C) FA(20:3), (D) FA(20:1), (E) FA(22:6), (F) FA(22:5), and (G) FA(22:4). No significant differences were observed between penumbra and core for these fatty acid species. Data are presented as mean ± SD. N = 3.

**Supplementary Figure S2.**
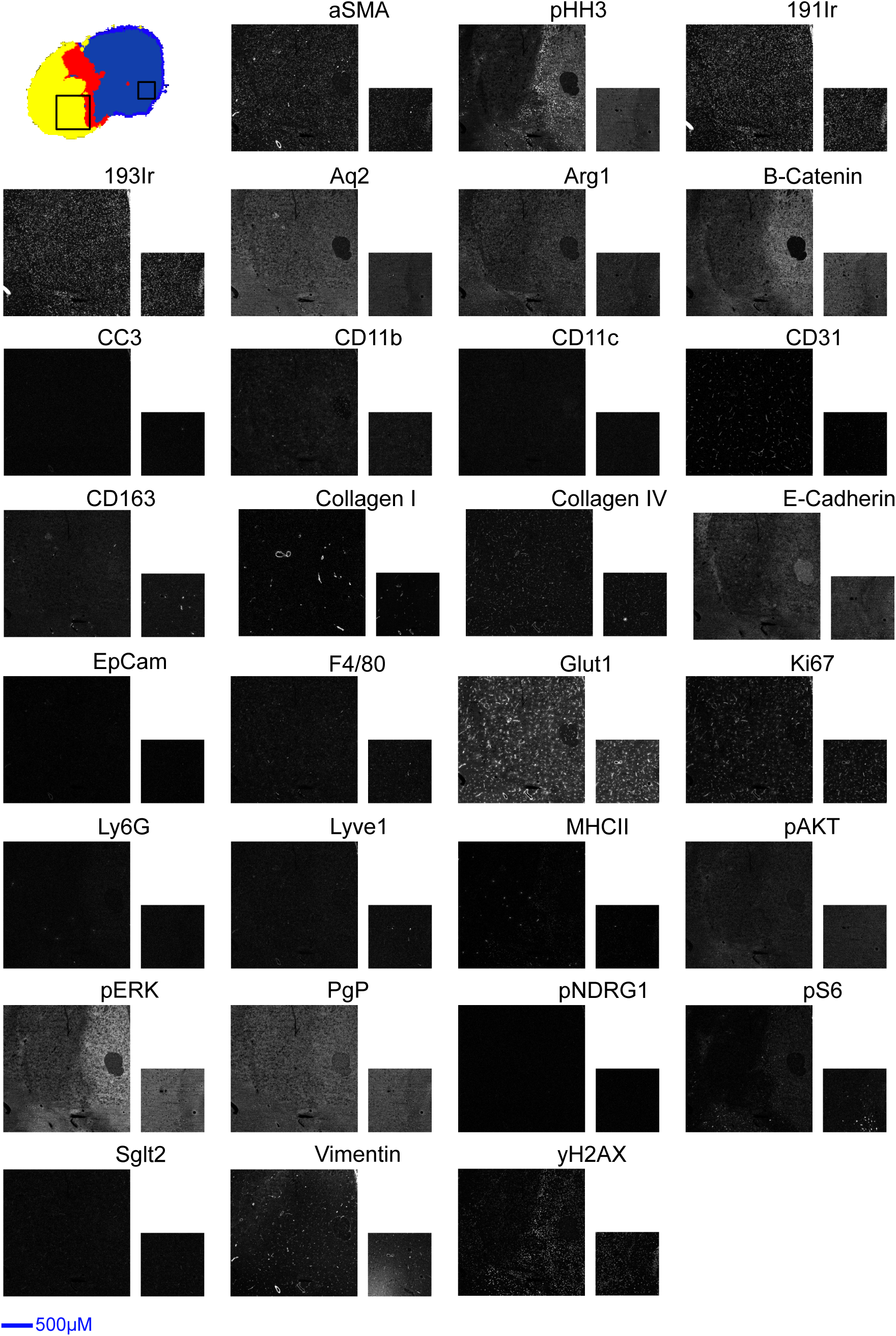
| Imaging mass cytometry profiling of healthy tissue, penumbra and ischemic core. The MSI-derived segmentation map (top left) with boxed regions indicating regions of interests for IMC. Larger boxes indicate ROI in ischemic hemisphere and smaller box represent ROI in the healthy hemisphere. Scale bar = 500µM.

